# AREAdata: a worldwide climate dataset averaged across spatial units at different scales through time

**DOI:** 10.1101/2021.10.05.463057

**Authors:** Thomas P. Smith, Michael Stemkovski, Austin Koontz, William D. Pearse

## Abstract

In an era of increasingly cross-discipline collaborative science, it is imperative to produce data resources which can be quickly and easily utilised by non-specialists. In particular, climate data often require heavy processing before they can be used for analyses. Here we describe AREAdata, a free-to-use online global climate dataset, pre-processed to provide the averages of various climate variables across differing administrative units (*e*.*g*., countries, states). These are daily estimates, based on the Copernicus Climate Data Store’s ERA-5 data, regularly updated to the near-present and provided as direct downloads from our website (https://pearselab.github.io/areadata/). The daily climate estimates from AREAdata are consistent with other openly available data, but at much finer-grained spatial and temporal scales than available elsewhere. AREAdata complements the existing suite of climate resources by providing these data in a form more readily usable by researchers unfamiliar with GIS data-processing methods, and we anticipate these resources being of particular use to environmental and epidemiological researchers.

## Background & Summary

High quality open-source weather and climate datasets are essential for researchers across the whole spectrum of scientific fields. Various sources of these data exist, such as the Copernicus Climate Data Store (CDS)^1^, Berkeley Earth^2^, WorldClim^3^, CHELSA^4^, and NASA’s Earth Data (https://earthdata.nasa.gov/). These data are generally available in a gridded (raster) format, with each climate variable estimated for a grid of latitudes and longitudes at a given spatial scale. In order to use these data, extensive processing is often required to extract climate variables for the region(s) of interest.

To aid researchers, static datasets of climate variables averaged across spatial units (*e*.*g*., countries) over a particular time period have been released (*e*.*g*. CCKP^5^) based on these gridded datasets. Such data are useful for many applications, but they are rarely regularly updated for up-to-the-minute modelling (*e*.*g*., in the COVID-19 response), do not include comparable forecast data (*e*.*g*., for climate change prediction), and are in spatial units not optimised for social science (*e*.*g*., countries or regions). In particular, the COVID-19 pandemic has demonstrated a need for easily accessible up-to-date climate estimates, as researchers have scrambled to assemble datasets to investigate the potential seasonality of SARS-CoV-2. The need to process these data leads to a duplication of effort as many researchers used GIS methods to quantify climate parameters across the same spatial units, using the same original data sources. This is particularly clear when considering the number of studies which have independently processed the same CDS ERA-5 data across similar (although often the same) spatial units to investigate the effects of environment on SARS-CoV-2 transmission^6–12^. Such duplication of effort is not just a barrier to reproducibility, but also to the efficient allocation of resources (particularly in the face of urgent crises like COVID-19).

We developed a pipeline to aggregate state-level climate variables in the USA across time as part of our work investigating SARS-CoV-2 seasonality^9^. Here, we extend this pipeline to cover additional climate variables and spatial units, which we release as the open-source resource AREAdata: “**A**dministrative **R**egion **E**nvironmental **A**verages data”. AREAdata provides global daily estimates of temperature, specific humidity, relative humidity, ultraviolet (UV) radiation, and precipitation at three different levels of spatial organisation, equivalent to countries, states and counties, from the beginning of 2020 to the near-present. Additionally, AREAdata provides population densities for the same spatial units, as well as an array of forecasted future annual mean temperatures from nine global climate models under four shared socio-economic pathway scenarios, based on the CMIP6 future climate projections^13^. These are provided as downloadable ‘RDS’ files for direct use in the R statistical programming language^14^, and as compressed tab-delimited text files for other applications.

## Methods

To produce the daily climate estimates provided in AREAdata, we gather gridded rasters describing daily climate data and average these climate variables across the geographic areas of spatial units at different levels of administrative organisation.

Below, all software packages given in *italics* are *R* packages (version 4.1.0)^14^ unless otherwise specified. The code to fully reproduce this pipeline is freely available under a GPL v3.0 license and can be acquired from our *GitHub* repository (https://github.com/pearselab/areadata). Continual updates of the output files as new climate data becomes available can be found on our *GitHub* project website (https://pearselab.github.io/areadata/). These continual updates are automatically released monthly, however the underlying code to run these updates locally is also shared so that users can update these data to-the-day when necessary. Output files for the county-level estimates are large (*>*100MB), and so are released on figshare (https://doi.org/10.6084/m9.figshare.16587311). Data on either platform are version-controlled with dates of submission recorded and past versions archived. To ensure that those who process and release the raw data going into AREAdata are properly acknowledged, a condition of use of AREAdata is the citation of the raw data, and this information is provided on the website.

### Data collection

We acquire shapefiles for worldwide administrative areas from the Global Administrative Areas (GADM) database^15^ at three different spatial scales: GID 0, GID 1, and GID 2. GID 0 is equivalent to countries, and (in the USA) GID 1 and GID 2 are equivalent to states and counties respectively.

We collect hourly estimates of climatic variables for the ERA-5 reanalysis from the Coperincus Climate Change Service’s Climate Data Store (CDS). Temperature (K), specific humidity (kg kg^−1^; mass of water vapour per kilogram of moist air), and relative humidity (%; water vapour pressure as a percentage of the air saturatation value) are acquired from the pressure-levels dataset^16^ at 1000 hPa (*i*.*e*., surface atmospheric pressure). Estimates of ultraviolet (UV) levels (J m^−2^; the amount of UV radiation reaching the surface) and precipitation (m; total precipitation, the accumulated liquid and frozen water falling to the Earth’s surface as measured in metres of water equivalent) are acquired from the surface-level dataset^17^.

Global population density data are acquired from the Gridded Population of the World collection, version 4, revision 11^18^. These data consist of population density estimates based on national and sub-national censuses and population registers. They use a gridding algorithm to assign population densities to grid cells, and these data are provided as rasters at different scales. Here we use the 15 arc-minute resolution for consistency with the resolution of the ERA5 climate data.

Downscaled CMIP6 future climate projections are acquired from WorldClim^3^. CIMP6 is the 6th phase of a global climate model (GCM) inter-comparison project, coordinating the design and distribution of global climate model simulations^13^. These model simulations are typically numerically complex and thus to facilitate fast computation, the world is divided into coarse grid cells. This is not ideal for studies investigating phenomena at higher spatial scales, and thus WorldClim provides downscaled versions of future predictions from GCM outputs, at higher spatial resolutions, based on WorldClim v2.1 as baseline climate. WorldClim provides these downscaled data for nine GCMs: BCC-CSM2-MR, CNRM-CM6-1, CNRM-ESM2-1, CanESM5, GFDL-ESM4, IPSL-CM6A-LR, MIROC-ES2L, MIROC6, MRI-ESM2-0, and for four Shared Socio-economic Pathways (SSPs): 126, 245, 370 and 585.

### Climate averaging pipeline

We use the Climate Data Operators program^19^ to compute daily means from the hourly data for each of the climate variables acquired from the CDS. We then calculate the mean value of each environmental variable across the administrative units given in each of our acquired shapefiles (*i*.*e*. countries, states, etc.), using the *exactextractr* R package. Specifically, we compute the mean of all grid cells fully or partially covered by the administrative unit polygon, weighted by the fraction of each cell covered by the polygon. When new climate data becomes available, these are appended to the previously extracted data to produce a single, live, updated output file for each administrative level and environmental variable combination. The data produced are simple files containing the daily climate estimates by spatial unit, *e*.*g*. country and by date, which we output as .RDS files for use in R and as zipped tab-delimited text files for other applications. We use an automated pipeline to produce new estimates on a monthly basis, which updates these files and automatically publishes new versions to *GitHub* and figshare (the links for which remain constant). An example of the input data for temperature from the 1st January, 2020, and the output following application of our methodology for country-level spatial units is given in Figure 1.

**Figure 1.**
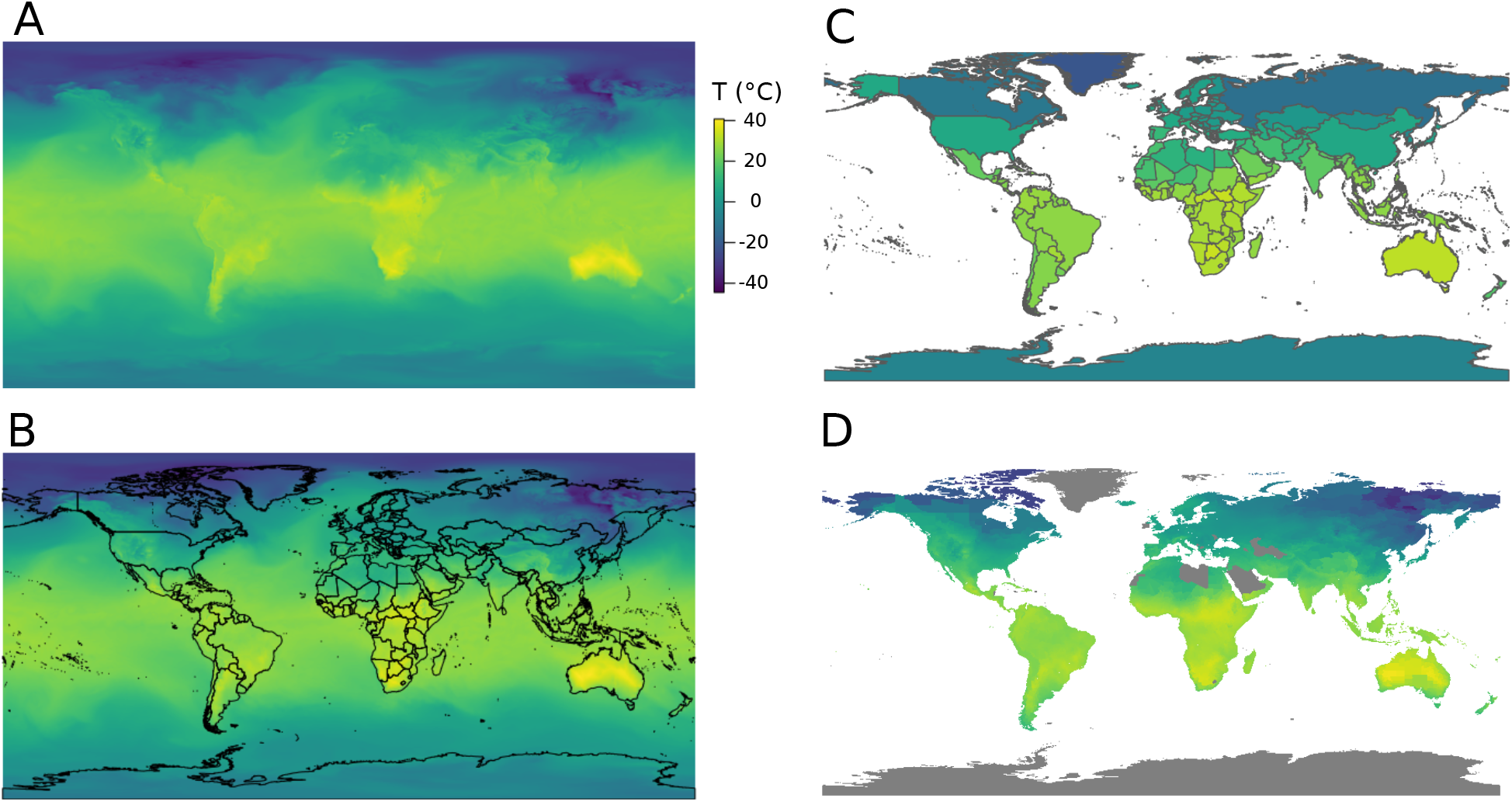
An overview of the processing pipeline of AREAdata and its resulting output. (A) Climate data rasters are acquired. The example figure shows daily mean temperature (°C) for 2020-01-01. These are hourly estimates that are averaged to daily estimates at a spatial resolution of 0.25×0.25°. (B) Shapefiles from GADM are used to demarcate the boundaries of different administrative units, in this case countries (GID 0). (C) The climate variable is averaged (mean) across each spatial unit. This averaging is weighted according to the relative amount of each grid cell within a region, which is of particular importance for smaller units (*e*.*g*., GADM level 2 regions). (D) This is the result of applying the same methodology to county-level (GID 2) regions. Here, for visualisation, land areas with no GID 2 demarcations are shown in grey, and border lines are not shown. The output more closely resembles the original data (panel A) at this finer scale. The data are released in .RDS and zipped .txt formats every month.

We use the same methods to process the gridded population density data, which we provide similarly with a single population density estimate for each spatial unit. We process annual mean temperatures from the climate forecast data, and again provide estimates by spatial unit for each combination of GCM and SSP. The population density and temperature forecast output files are static (not continually updated). Our website provides an easy interface to download these data; however, users can also run the provided code locally to make adjustments to the calculations and generate their own files.

## Data Records

AREAdata can be accessed at our *GitHub* site (https://pearselab.github.io/areadata), which contains download links to each data file.

These are distributed both as .RDS files for use in the R statistical programming environment and as zipped tab-delimited files for other uses. The daily climate files consist of a matrix of point estimates of an environmental variable (either temperature, specific humidity, relative humidity, UV or precipitation), with rows representing each spatial unit that the variable was averaged across and columns representing the date. The population density files consist of a matrix with a single column of population density point estimates, with rows for each spatial unit. The climate forecast files consist of a matrix of point estimates for annual mean temperatures, with rows representing each spatial unit, and columns representing the combination of global climate model (GCM) and shared socio-economic pathway (SSP), and the year range of the projection. Column headers for the forecasting files follow the labelling convention <GCM>_<SSP>_<XXXX-YYYY>, where XXXX-YYYY specifies the date range of the forecast. These files are all distributed by the level of spatial organisation that the data have been averaged across (*i*.*e*. separate files for countries, states, counties). In the initial release, AREAdata provided daily climate estimates from 2020-01-01 to 2021-08-31.

A visualisation of the daily temperature estimates for different levels of spatial organisation is shown in Figure 2.

**Figure 2.**
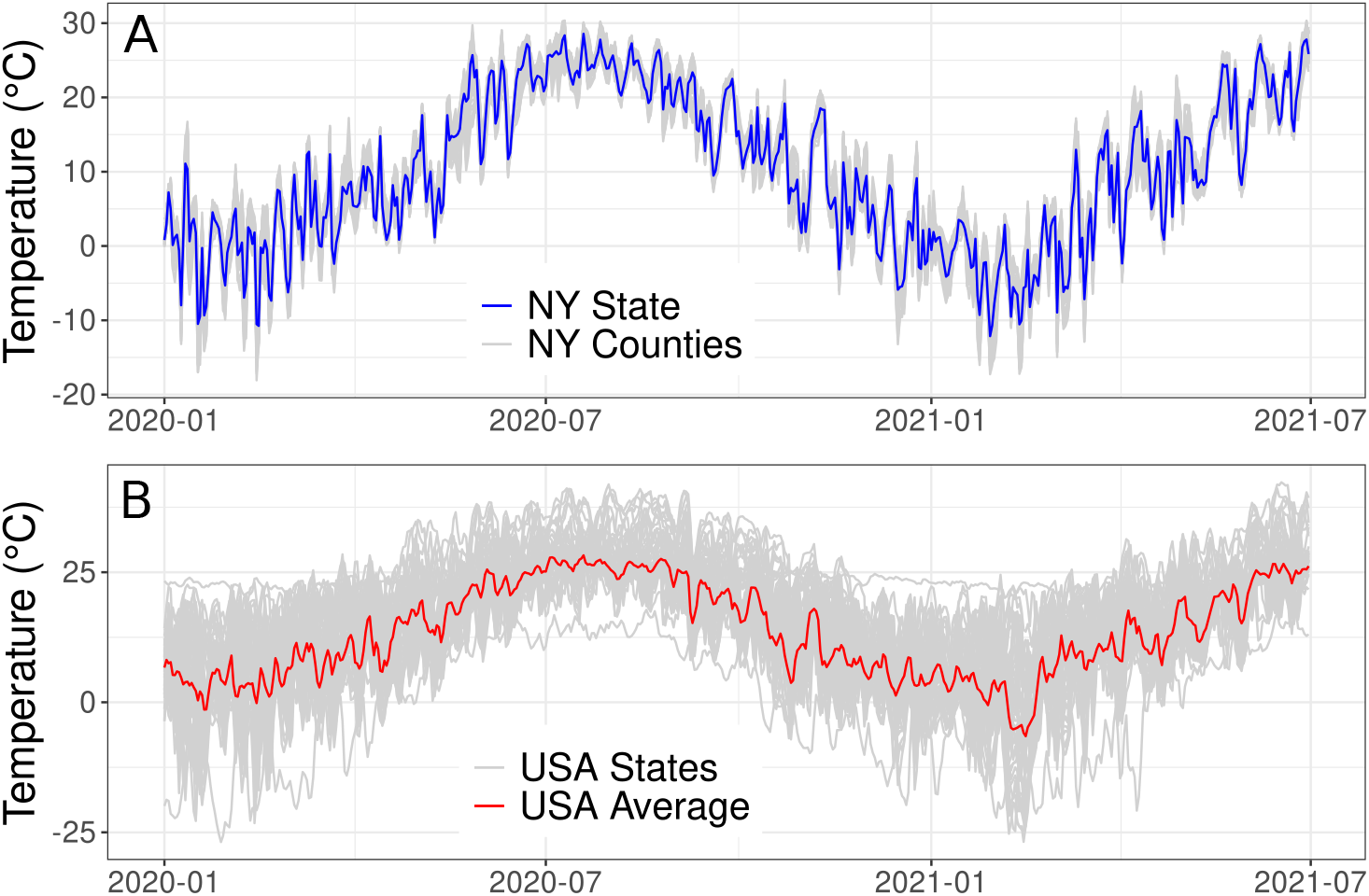
Visualisation of output daily temperature estimates for the USA, spanning 18 months from 2020-01-01 to 2021-06-30. (A). Grey lines are the county-level (GID 2) estimates for each of the 63 counties in New York State, blue line shows the state-level (GID 1) estimates for New York State. (B). Grey lines show the state-level (GID 1) estimates for each of the 51 states of the USA (50 states plus Washington DC), red line shows the country-level (GID 0) estimate across the whole of the USA. As expected, there is much greater variation in temperature between states (B) than between counties within the same state (A).

## Technical Validation

All of the underlying daily climate data contributing to the main product of AREAdata were acquired from the CDS ERA-5 reanalysis. The reanalysis is a data assimilation technique which combines a wide range of meteorological observations with a prediction model, to produce an improved gridded observational dataset^20^. The benefits of using the ERA-5 reanalysis in particular include its frequent updates (five days into the past) and high temporal resolution (1hr)^20^. The data underlying the daily climate estimates in AREAdata have thus been heavily validated and are already widely used, *e*.*g*.^7,8^.

The CDS ERA-5 climate data is available as gridded values at the 15 arc-minute (0.25°) scale, equating to roughly 27×27 km grid cells at the equator. Very few countries have areas smaller than this, however here we investigate whether these data are granular enough for use with the smaller spatial units. When looking at the distribution of unique grid cells spanned by each spatial unit (*i*.*e*., the number of climate values that the spatial polygon covers and is thus averaged across) we find that the vast majority of countries and states span many grid cells (Figure 3). However, the county-level (GID 2) administrative units are generally much smaller with most spanning fewer than 10 unique grid cells, including the 10% of counties represented by only one climate estimate (Figure 3). This may lead to climate estimates being duplicated for the smallest areas if there are instances where two or more areas are entirely encompassed by the same grid cell, thus returning the same climate estimate. To address this, we tested whether any regions return the same climate value through time. We found no instances of this duplication within our datasets – each GID 2 region has a unique climate profile. We therefore conclude that the underlying data, although somewhat coarse, are granular enough to return distinct climate estimates at the GID-2 scale. The code to run this analysis is available in our *GitHub* repository.

**Figure 3.**
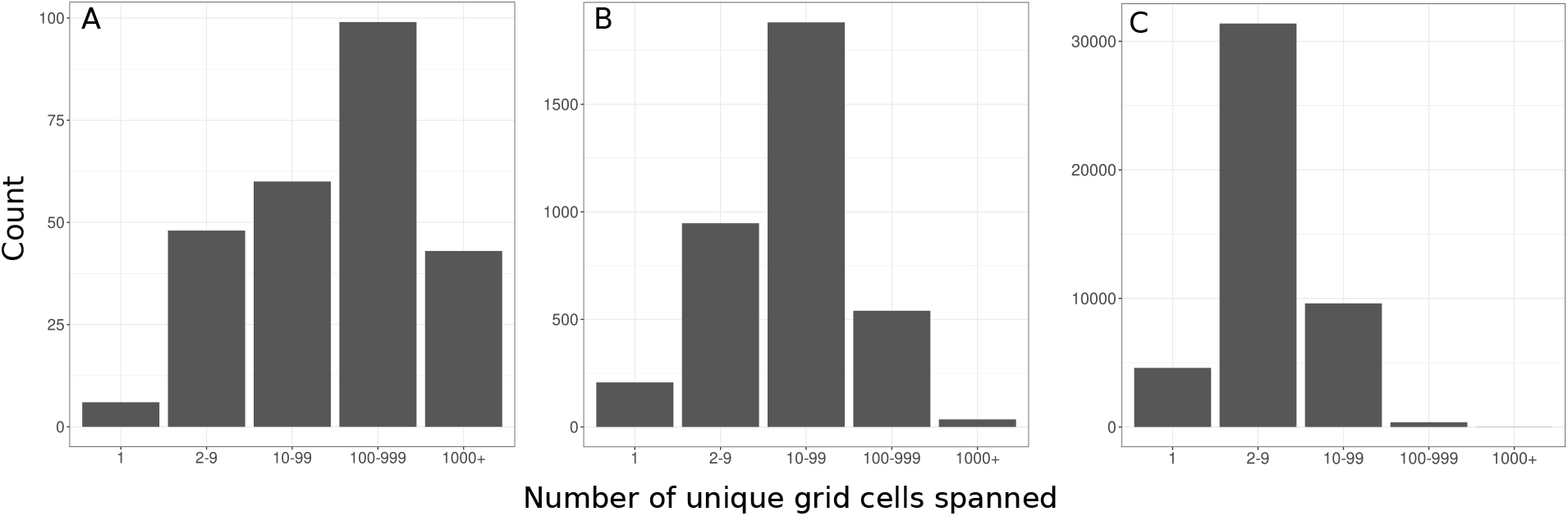
Variation in the number of distinct grid cells fully or partially covered by spatial area polygons across scales in (A) countries (GID 0), (B) states (GID 1), and (C) counties (GID 2). Across all scales, most administrative areas are large enough that their climate estimates are based on multiple grid cells of the climate rasters. This shows that the underlying data are largely granular enough for our usage; however, we further tested this by ensuring regions have distinct climate profiles (see Technical Validation).

Few publicly available datasets exist to provide climate estimates for spatial regions through time to validate our daily outputs against. However, the Climate Change Knowledge Portal (CCKP) maintains an open dataset of historical monthly climate estimates at the country level, which includes mean temperature^5^. The CCKP data is not based on the CDS ERA-5 climate reanalysis, but derived from the “Climatic Research Unit gridded Time Series” (CRU-TS) dataset; a monthly gridded climate dataset at a 0.5° latitude/longitude scale^21^. Although these climate estimates are derived from a more coarse underlying dataset than AREAdata, both spatially (0.5° rather than our 0.25° grid) and temporally (monthly, as opposed to our daily estimates), this does provide a potentially useful comparison. Here, we aggregate our daily temperature estimates for countries to monthly means, and compare these to the CCKP estimates for 2020. When comparing monthly estimates of mean temperature for each month in 2020 from 182 countries within the CCKP data and AREAdata outputs (*i*.*e*. 2,184 point estimates), we find an overall Pearson’s correlation coefficient of 0.935. There are systematic differences in the AREAdata and CCKP outputs, with AREAdata consistently producing marginally higher temperature estimates than the CCKP estimates, especially in lower temperature countries (Figure 4). The CRU-TS dataset (upon which the CCKP estimates are based) is observational, whereas the ERA-5 data (upon which AREAdata is based) is a reanalysis combining forecasting model estimates with observations to provide spatially and temporally finer-grained climate estimates. Previous work has found similar overall correlations between the CRU-TS and ERA-5 data (approx 0.9)^22^, however these datasets can vary significantly at particular times and in particular places^22,23^. In particular, there is likely a warm bias in the ERA-5 reanalysis in recent years due to lesser representation of snow on top of sea ice^24^, which may explain the deviations we observe here. Therefore, despite the expected deviations due to differences in underlying datasets, the strong correlations we observe between AREAdata and the CCKP validates our methodology by showing that our country-level estimates are largely consistent with another source of country-level climate estimates through time.

**Figure 4.**
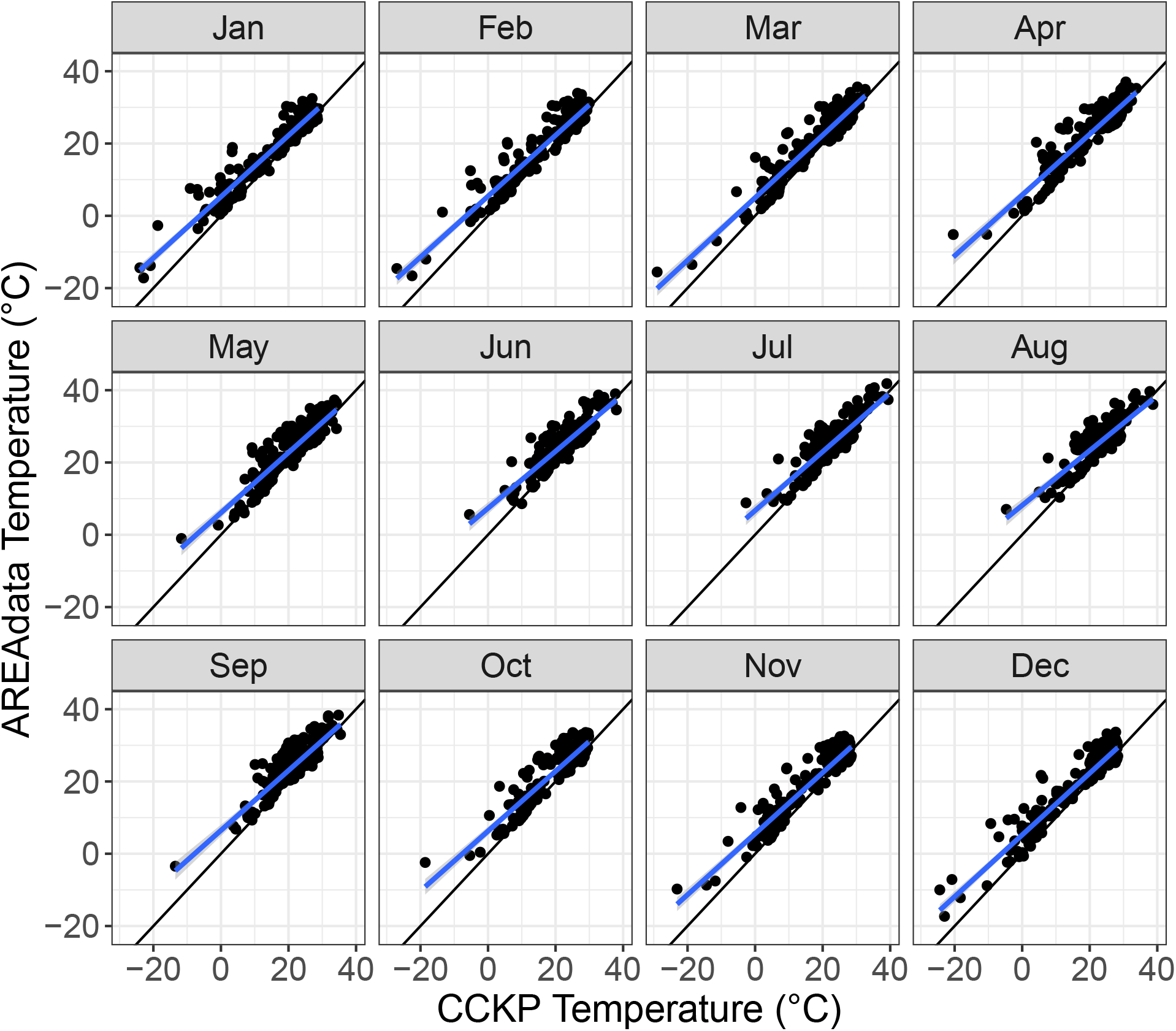
Comparison between country-level monthly mean temperature estimates obtained from the Climate Change Knowledge Portal (CCKP)^5^, versus the daily estimates from AREAdata aggregated into monthly means. The CCKP estimates are derived not from the CDS ERA-5 data, but from the Climatic Research Unit gridded Time Series (CRU-TS). Despite the differences in underlying data and methods, we find strong correlations between AREAdata and the CCKP temperature estimates (all *r >* 0.84). This validates our work by showing that our methodology produces sensible temperature estimates in line with an alternative dataset.

To ensure validity of the downloaded climate estimate files, we provide a file with MD5 checksum values with each monthly data release. This is a hash function computed on a file, and if two files have the same value then it is very likely that they are identical since a single change to a file will radically alter its checksum. A script to automatically check the published MD5 checksum values against the downloaded files and note any discrepancies is provided in our *GitHub* repository.

## Usage Notes

The AREAdata point estimates are distributed as .RDS files for use in the R statistical programming language, which can be imported with the readRDS() command. For example, to import daily temperature data for countries from the main *GitHub* directory after cloning the repository: readRDS(“output/temp-dailymean-countries-cleaned.RDS”). Alternatively, the data could be loaded directly ’live’ from the website itself: readRDS(“https://github.com/pearselab/areadata/raw/main/output/temp-dailymean-countries-cleaned.RDS“), for example as part of a script that made use of the latest daily temperatures for near-real-time modelling and forecasting. The data objects are two-dimensional matrices with rows corresponding to administrative regions and columns corresponding to dates. The values of the matrix are in the units of the environmental variable in the given file. Each administrative region is described by the codes used by the GADM, however we also supply a metadata file linking these codes to place names (“data/name-matching.csv”). Place names from this metadata file could be added to the AREAdata objects in R by transforming them into dataframes and then using the merge() function.

We also distribute these data as zipped tab-delimited .txt files for use in other applications. However, these could also be directly read into R in a single line if required, *e*.*g*. read.table(unz(“output/temp-dailymean-countries-cleaned.zip”, “temp-dailymean-countries-cleaned.txt”), check.names = FALSE, row.names = 1), or using a URL (as in the example above) if desired. Note that the zip compression is less efficient than the RDS compression, resulting in larger files, so for R applications the .RDS files are recommended.

## Code availability

All code underlying AREAdata is available from our team’s *GitHub* repository (https://github.com/pearselab/areadata).

## Acknowledgements

This work was funded by Natural Environment Research Council Grant NE/V009710/1.

## Author contributions statement

WDP and TPS conceived the work, WDP, TPS and MS conducted preliminary work, TPS and AK tested the code, TPS finalised the results, and TPS wrote the manuscript. All authors reviewed and edited the manuscript.

